# Cerclage Wire as an Affordable Alternative for Internal Fixation in Murine Critical Sized Defect Models

**DOI:** 10.64898/2026.07.24.740565

**Authors:** Kaitlyn Cimney, Shivani Sawant, Kell Sprangel, Zachary Osborn-King, Khady Diop, Jordin Marshall, Allison Smith, Adam Kling, Devin Medved, Colton Sterling, Kevin Wolf, Ryan Brune, Gennadiy A. Busel, Angela C. Collins, Daemeon Nicolaou, Benjamin A Walter, Sara McBride-Gagyi

## Abstract

Critical sized defects (CSDs) are a serious challenge in orthopedics that require the development of more robust and effective treatments to improve quality of life for patients. Current CSD research is limited by the applicable and affordable animal models available. Mice would be the preferred species as they are cheaply housed and have many transgenic variations readily available; however, their small size makes CSD surgeries difficult and expensive. We propose the use of cerclage wires to achieve internal plate fixation. PEEK plates were secured to the right femur of 26 C57BL/6 mice using four cerclage wires to achieve modified double-loop fixation implemented through bicortical holes and defects were created. 10 received 3mm defects and 6 received 4mm defects that were left empty and were taken out to 20 weeks (Group E3, E4). Another 10 received 3mm defects that were filled with a morselized bone graft and were taken out to 8 weeks (Group G3). Blinded longitudinal x-ray grading by orthopedic surgeons was conducted on the empty defects for plate stability and wire fixation. All samples received microCT analysis at their endpoints. There were no significant differences in plate or wire stability between the empty groups and wire scores worsened negligibly over time. MicroCT analysis further supported wire integration as bone growth directly upon the wires was observed in all samples. The efficacy of this model in achieving non-union when left untreated was also confirmed via microCT. Further, only three mice in group G3 achieved union and two of these unions were not optimal. Our study is the first to successfully show that cerclage wire can be used in a murine CSD model to achieve affordability and clinical relevancy.

## Introduction

Critical sized defects (CSDs) pose a serious challenge in orthopedics and negatively affect physical and financial quality of life for millions globally [1]. CSDs are bone defects of at least 2.5 cm in humans, or generally twice the diameter of the affected bone, and are unable to heal spontaneously without surgical intervention. If a CSD cannot be reconstructed, the bone cannot serve its intended weightbearing, locomotive, or protective functions. While not naturally occurring, CSDs occur clinically due to high-energy injuries (e.g. car crash, blast injury), infection, and osteosarcoma resection [2,3]. Limb salvage, rather than amputation, is desirable whenever possible due to the long-term benefits of increased patient quality of life and lower life-time costs. Amputation is less expensive and quicker in the short-term but remains a significant physical and fiscal burden for the rest of the patient’s life. More modern techniques like distraction osteogenesis (DO), traditional bone grafting, and Masquelet’s induced membrane technique (MIMT) have enabled limb salvation with high success (i.e. ∼95%, ∼90%, and ∼86-90%, respectively) but are still not ideal [4–14]. DO employs an external fixator that can be cumbersome and painful. Also, treatment effectiveness is highly dependent on patient compliance and regular clinical follow up. Traditional bone grafts require less maintenance but have limited applicability in larger defects (>5cm). MIMT has been shown to have extraordinary healing abilities (>25cm), but very little is known about its mechanism of action, making improvements and repeatability difficult [4,7,13]. Thus, further development of CSD treatments such as novel grafts, graft extenders, and graft additives, as well as studying biological mechanisms of successful traditional grafting and MIMT-facilitated grafting, are driving factors of this project.

CSD research is underpopulated in part due to the financial barriers associated with animal models. Decades of medical research has established rodents are useful models for replicating human physiology as their biological mechanisms are often very similar. This is especially true in orthopedics where the composition, structure, and biology of growth and repair of rodent bones closely resembles that of humans [15–17]. Additionally, rodents offer cheaper housing options than larger species like sheep or pigs [16]. Most CSD research is performed in rats as their larger size makes the complex surgical operations much easier, but the lack of readily available transgenic rats limits the depth of mechanistic studies [15–17]. The use of transgenic mice allows us to address bigger questions in CSD research, especially with the emergence of tissue-engineered treatments and the MIMT mechanism. However, mice are about 1/10^th^ the size of a rat, making their surgeries technically complex due to the dimensions and financially inaccessible due to the necessary micro- scale implants and high-precision tools [17].

Clinically, fixation can be achieved either internally (i.e. plate or intramedullary rod) or externally, with internal fixation usually being the preferred [18,19]. Using animal models requires extra care to ensure the animal doesn’t disturb the implant or change how it walks, which could lead to experimental failure [20–22]. While both internal methods are extremely clinically relevant, intramedullary rods are not ideal for use in rodent CSDs because implanting bicortical transverse interlocking screws to prevent axial translation and rotation is particularly challenging [23]. In contrast, plate interlocking fastener implantation is much easier, making plate fixation the preferred and more stable method. Further, the implant’s dimensions are not as limited [20]. So, mechanically adequate plates can be fabricated from many different materials which enable radiolucent material use (e.g. polyether ether ketone (PEEK)).

Despite internal plates’ advantages, screws or pins are still needed. Mouse screws have to be extremely small because mouse femora are only ∼1.5 mm in diameter. Micro-screws are commercially available from custom or bulk manufacturers, but their costs can be well over $50.00 per mouse, exponentially driving up costs and making murine CSD models price prohibitive [20–25]. Another fixation method used clinically is cerclage wiring, which is much more economical in both the clinical and laboratory setting. Cerclage wire can be made of either metal or plastic, with metal being the most common, and can be single or multi-stranded (e.g. braided or cabled). Cabled wires offer more rigid fixation while braided wires have been shown to fail much earlier [26]. There are four categories of wire use in fixation: tension band, cerclage, hemi-cerclage, and interfragmentary wires. We will be focusing on the cerclage method which involves wrapping the wire around the entire bone. Cerclage wiring can be used as either the primary or secondary form of fixation, with the latter being the more common in humans as the wires are often not strong enough to withstand the internal forces associated with fracture recovery. In fact, it is currently experiencing a revitalization for use in preventing/managing periprosthetic fractures, especially in the femur [26–28]. In small animals, however, cerclage wires do not undergo massive forces, potentially allowing for their use as the primary fixation technique [27,28]. Furthermore, estimated cost per mouse is $0.08 when using commercially available 36G stainless steel wires, offering fixation at a cost of at most 0.2% of commercially available microscrews.

The main concern when using cerclage wire, especially as the primary fixation, is breakage and/or loosening due to its poor resistance to interfragmentary forces. This can lead to further complications which impair healing, including compromising the blood supply via strangulation and bone resorption/lysis due to excessive implant movement [26–28]. Mitigation of these concerns can often depend on the method of wire fastening. There are two main steps involved in fastening: 1) securing the wire, and 2) closing. Twist, single-, or double-loop knots can be used to secure either a single or multi-stranded wire. Twist knots are formed by twisting the ends of the strand while pulling. Looping simply refers to the number of times the wire is wrapped around the entire bone. Single-looping involves creating a loop on one end of the strand, wrapping the single strand around the entire bone, and then passing through the created loop. Double-looping is achieved by folding the wire in half and wrapping the two strands around the bone and back through the loop in parallel [27]. Double-looping is preferred as it offers more stability to the implant; this is especially true if using single stranded wire as it has been shown to have comparable mechanical strength to a single-looped multi-stranded wire [26]. Closing the wire can be accomplished via twist, crimp, or various knot configurations [28,29]. Twisting the wire ends is the most popular method used clinically for its simplicity and effectiveness. The knot twist is also a preferred method which employs a simple knot following by twisting, as well as the “secured twist” which involves a simple twist that is then folded over to prevent unwinding [28,30]. To our knowledge, this has never been attempted in mice and has only been reported twice in rats. These companion rat studies used a simplified technique of four cerclage wires wrapping around both the plate and whole bone without medial-lateral stability [31,32].

Due to its clinical use in humans and animals, cerclage wire was pursued as a more economical alternative to microscrews at a fraction of the cost. The objective of this study was to develop a cost-effective murine CSD model which will not heal even when grafted. We hypothesize that the proposed modified double-loop fixation implemented through bicortical holes will offer sufficient stability to preserve the critical-sized defect while minimizing iatrogenic damage to the bone, ensuring the model remains a true non-union even after osteogenic grafting intervention.

## Methods

### Plate Material Selection & Fabrication

Polyether ether ketone (PEEK) was chosen as the plate material due to its excellent mechanical properties, biocompatibility, and radiolucency as well as standard use in rodent bone repair studies [33,34]. PEEK plates were laser cut from a 1.5 mm thick sheet of PEEK to final over all dimensions of approximately 2 mm x 10.5 mm. The plates’ detailed geometry is available in the Supplement (**Error! Reference source not found.**). A slightly different plate geometry 3D printed from tough 1500 (T1500, FormLabs) was also tested. However, this material was found to be exceptionally unstable and weak *in vivo* resulting in early removal from the study for most animals. Thus, for animal welfare concerns, T1500 was abandoned as a viable plate material and further study halted. The data collected for these animals is provided in the supplemental section to underscore the findings presented in the main manuscript.

### Experimental Animals and Groups

All animal procedures were approved by the IACUC committee at The Ohio State University (Protocol #2022A00000082). Male and female C57BL/6 wild type mice were assigned to one of three treatment groups defined by the defect size and whether morselized bone graft material was added (Table 1**Error! Reference source not found.**). The empty 3 mm defect group (E3) represents the ideal scenario while the empty 4mm defect group (E4) represents a challenge scenario with a larger CSD. Finally, the grafted 3 mm defect group (G3) represents an osteogenic grafting intervention.

**Table 1.** Group Demographics.

| Group | Scenario Objective | Defect Size (mm) | Graft Status | Number of Mice (Male/Female) | Age (weeks) |
| --- | --- | --- | --- | --- | --- |
| E3 | Ideal Fixation | 3 | - | 10 (4/6) | 16.5±0.62 |
| E4 | Challenge | 4 | - | 6 (5/1) | 16.4±0.29 |
| G3 | Clinical | 3 | + | 9 (3/6) | 16.4±0.36 |

### Surgical Protocol

Animals received buprenorphine extended release (1mg/kgBW) the morning of surgery. Each animal was anesthetized via isoflurane and prepped immediately before surgery by first applying eye ointment, removing hair with a depilatory cream, and then administering a local analgesic into the right hind limb (bupivacaine hydrochloride injection, <8mg/kgBW). The skin was aseptically prepped via 3 alternating passes each of chlorhexidine and isopropyl alcohol. The animal was then moved to the sterile surgical field and covered with sterile PressNSeal [35]. A small hole was cut in the PressNSeal near the right hind limb. The limb was brought through the hole and secured to a custom 3D-printed surgical stand to allow for maximal stability and clear access to the right femur.

The trochanter ridge of the right femur was located via skin palpation, and an incision was made in parallel to the femur from the trochanter to the knee. The skin was bluntly dissected from the muscle. The quadriceps and hamstrings were carefully separated and then bluntly dissected from the bone on all sides. A thin metal plate was inserted underneath the bone and over the muscle and skin to protect them during drilling. A custom 3D-printed drill guide was lined up with the trochanter ridge and secured to the bone using suture. Four 0.4 mm diameter pilot holes were drilled into the anterior-medial surface, after which the guide was removed. The plate was then attached to the bone by inserting one tail of a piece of 36-gauge stainless steel wire (Master Wire Supply, SKU: MWS-316L-36-0025-RO) into the proximal-most hole of the bone and plate in an anterior to posterior direction. The other wire tail was then inserted into the same hole to form a loop at the plate surface (Figure 1A). One wire tail was pulled to the femur’s medial side while the other was pulled around the lateral side (Figure 1B). The tails were threaded through the loop on top of the plate in opposing directions and tightened to form a self-fastening knot (Figure 1C-E). Before completely tying off, this was repeated for the distal most hole to ensure that all of the drilled holes lined up with the plate. To complete the knot, both tails were then looped around the bone again (Figure 1F-G), tied in a simple knot (Figure 1H-I), and twisted together on top of the plate (Figure 1J) to ensure security and remove any remaining slack. The ends were trimmed (Figure 1K), and the twisted wire was folded over the plate to limit tissue damage (Figure 1L). This process was repeated for the remaining holes. We recommend tying one of the distal holes second to prevent misalignment from overtightening at the proximal end. Additionally, the wires were wrapped slightly away from the area of the future defect on the middle proximal and middle distal holes to ensure that the wires would not slip into the defect. Once the plate was secured, a custom 3D-printed saw guide was slotted over the plate, and a defect was created using 0.13 mm Gigli wire. After removing the defect, the incision was either immediately closed (E3 and E4) or a morselized bone graft was implanted in the defect and then closed (G3) using interrupted stitches (surgeon’s knot, 5-0 monofilament suture).

**Figure 1.**
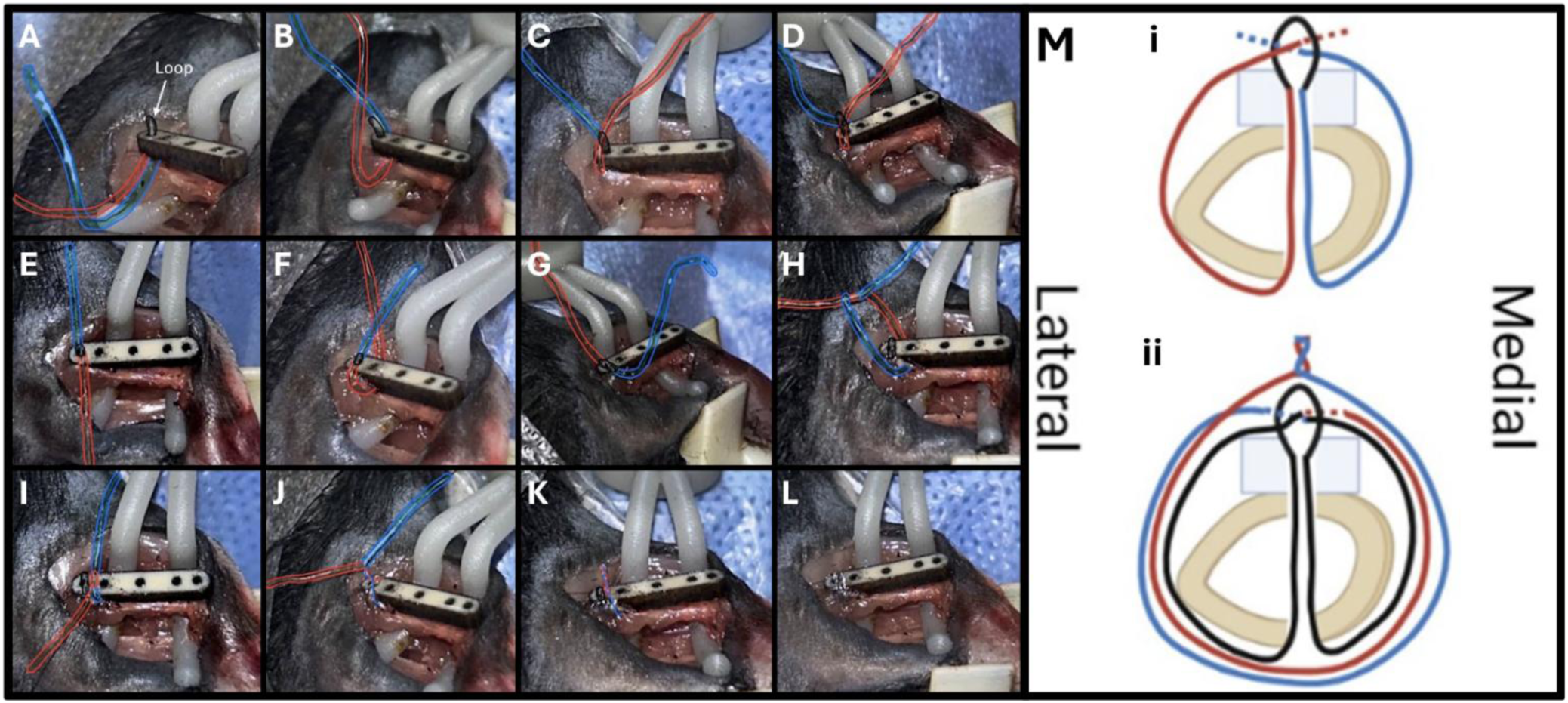
Wire Tying Procedure. **A.** A piece of wire is folded in half. Both ends are passed through the plate and then both cortices of the bone and pulled to form a small loop on top of the plate. **B.** One strand (blue) is passed under the bone to the medial side of the plate. **C.** The lateral wire is threaded through the loop in a proximal-to-distal fashion. **D.** The medial wire is threaded through the loop in a distal-to-proximal fashion. **E.** Both strands are pulled in opposite directions to tight the loop on top of the plate and form a self-fastening knot. **F.** The lateral wire is passed under the bone to the medial side. **G.** The medial wire is passed under the bone to the lateral side. **H.** The strands are used to make a simple knot. **I.** The simple knot is tightened to the plate surface. **J.** The strands are twisted together three times in a clockwise fashion. **K.** Excess wire is cut above the twists. **L.** The remaining wire is folded over. **M.** A graphic of the cross-sectional view of wire tying; **i** represents **A-E** while **ii** represents **F-K**.

**Figure 2.**
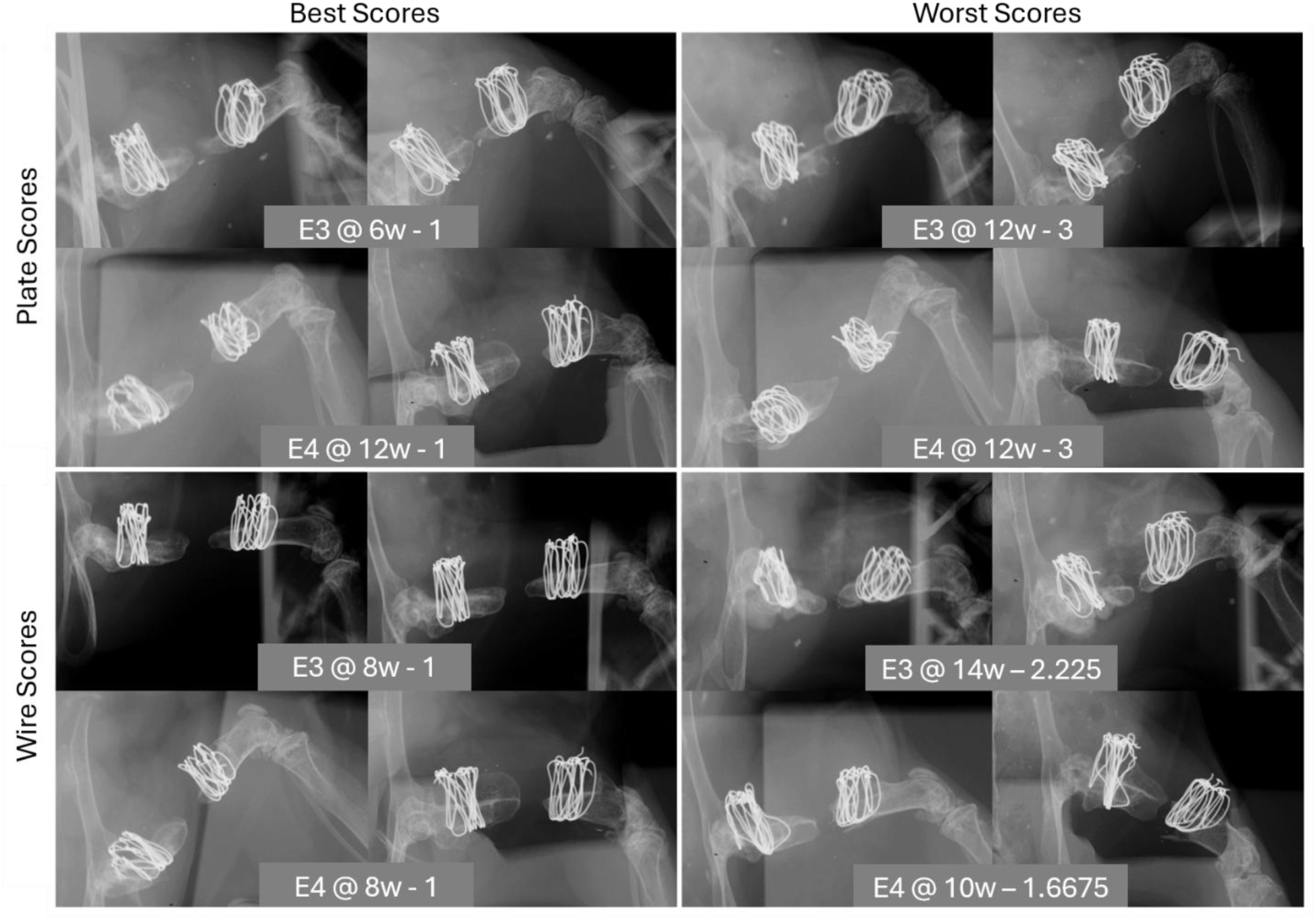
Representative x-ray images based on best (left) and worst (right) average scores for plates (top) and wires (bottom).

Bone grafts were harvested from sex-matched littermate donor animals euthanized via carbon dioxide exposure, followed by cervical dislocation. After euthanasia, an incision was made from the ankle to the hip and the skin was bluntly dissected from the muscle around the entire leg. The hip joint was located, and the tip of a scalpel was inserted to disarticulate. The knee was disarticulated by locating the anterior cruciate ligament (ACL) and inserting the tip of a scalpel between the distal end of the femur and proximal end of the tibia. The soft tissue was removed from the now free femur, followed by the removal of the femoral head and all tissue distal to the growth plate. The femur was morselized in a beaker using a rongeur until all large pieces were crushed and the graft was relatively homogeneous. Sterile PBS was then added to the beaker to preserve the integrity of the graft until implantation. The graft was left in this state for an average of 1 hour and 51 minutes ±48 minutes. Each donor produced graft material for 1 to 3 recipients.

### Longitudinal X-ray Grading

After surgery, each animal received anterior-posterior and medial-lateral x-rays biweekly until euthanasia. Groups E3 and E4 were imaged up to 20 weeks to qualitatively assess 1) the status of the plate and 2) the stability of the wires. Animals were humanely euthanized if a broken plate or adverse reaction (e.g. abrasions at incision site) was discovered (T1500 only). Group G3 were imaged over 8 weeks before euthanasia but only to monitor general post-surgical progression and for adverse events.

For the E3 and E4 animal images, unbiased orthopedic surgeons were given standard grading scales and asked to blindly assess randomized x-ray pairs (10 timepoints/animal; N = 160) based on plate alignment and wire integration (Supplemental Documents). To assess plate damage, the graders were told to compare the alignment of the distal bone segment with the proximal segment. If the segments were perfectly aligned, the x-rays are given a 1. A slight (2) or major (3) bend was indicated by an angle of 170° or 90° between the segments, respectively. A broken plate (4) was determined by looking between the views of the bone; if the distal segment rotated between the views while the proximal segment did not, the plate was deemed broken. To assess wire integration, the graders were told to look for bone lysis and overgrowth, both of which are reactions to unstable fixation. Lysis can be seen in the widening of the wire hole while overgrowth is evidenced by new bone formation over the ends of the wires in a volcano-like fashion. Integration was perfect (1) if neither reaction was present, fair (2) if either was present, and poor (3) if both were present. The graders’ scores were averaged for each individual animal and then averaged for each group.

### MicroCT Analysis

After euthanasia, all operated femurs were harvested from all groups and fixed overnight using 10% neutral buffered formalin and then kept in 70% ethanol until microCT scanning (MicroCT 50, ScanCo Medical; 70kVp, 114μA, 8W, 10μm). Images were processed using Dragonfly software (Version 2022.2 for Windows, Object Research Systems, Inc.). Quantitative analysis of bone repair included BV, TV, BV/TV, BMD, and TMD within the defect. Wire integration was observed qualitatively due to beam hardening from the cerclage wires. Scores were based on the presence or absence of bone growth directly upon cerclage wires within the drilled holes to assess integration. 3D reconstructions were used to determine union status.

### Statistics

Longitudinal x-ray grading was analyzed using a cumulative link mixed model (CLMM) with a logit link to assess plate and wire status over time within groups. Post Hoc pairwise comparisons were performed using estimated marginal means with Tukey adjustment where appropriate. This model was chosen to account for the ordinal nature of the grading scores and for repeated measurements within mice, anatomical wire regions, and grader variability. Fixed effects included group, week, anatomical wire region, sex, and the interactions group*week and group*wire region. Random intercepts were included for mouse, grader, and nested mouse-wire region combinations.

Quantitative microCT analysis was performed using one-way ANOVAs, followed by post hoc Tukey. Qualitative integration was assessed using Chi Squared Pearson Tests.

## Results

### Longitudinal X-ray Grading

Groups E3 and E4 were not significantly different in plate stability scores (p = 0.149) and changed similarly over time (p = 0.192), which slightly improved (p = 0.049) (Figure 3). Sex did not significantly influence plate stability (p = 0.280). For the wires, fixation slightly worsened over time (p < 0.0001), but there was no significant difference in the rate of deterioration between E3 and E4 (p = 0.455) (Figure 4). Anatomic regional differences were found both between and within groups. E3 had significantly worse fixation at the proximal most (p < 0.0001) and proximal middle (p = 0.017) wires versus E4 while there were no significant differences at the distal middle (p = 0.089) or distal most (p = 0.826) wires. Within the E3 group, the proximal most wire performed significantly worse than both the distal middle (p = 0.0235) and distal most (p = 0.0004) wires. Additionally, the proximal middle wire performed significantly worse versus the distal most wire in this group (p = 0.0089). There were no significant differences in anatomical region in the E4 group. Sex did not significantly influence fixation outcome (p = 0.465).

**Figure 3.**
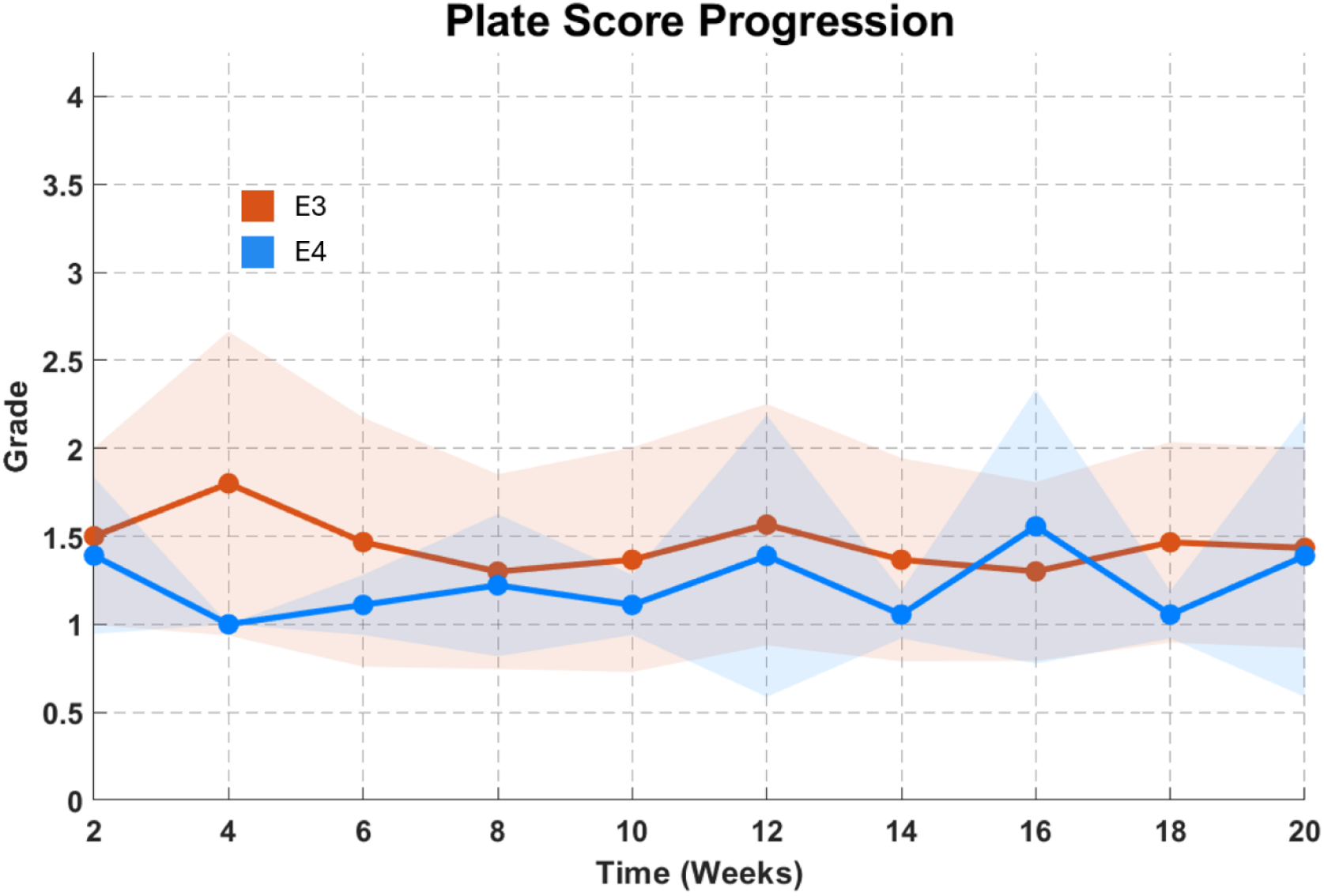
Plate score progression over time. Plate scores were similar between groups and relatively steady over the observation period.

**Figure 4.**
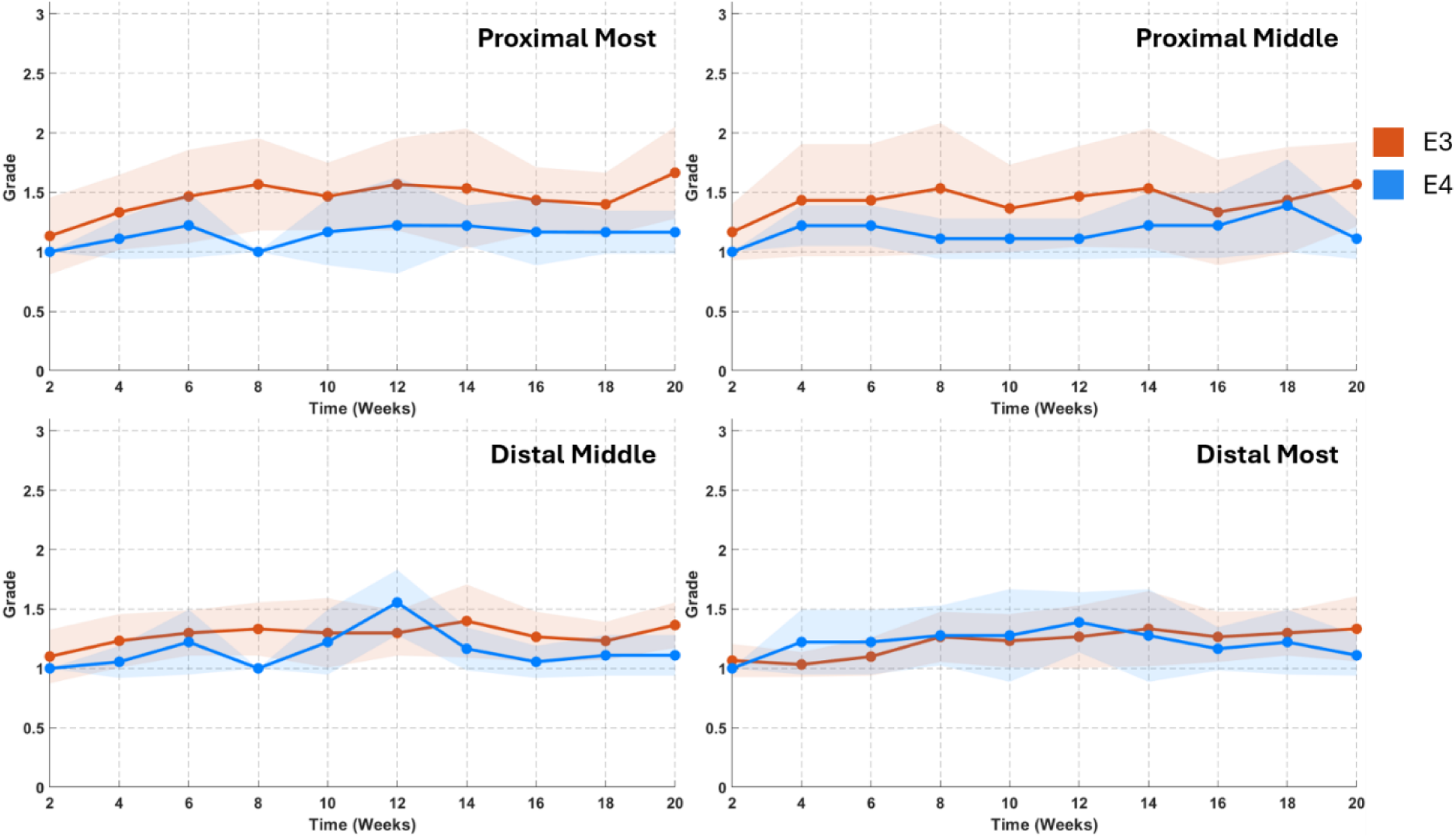
Wire score progression over time for each region. E3’s proximal most and proximal middle wire scores were overall significantly worse than E4’s while there were no differences for the distal wires. Scores statistically worsened over time, but overall rate of deterioration was not significant between groups and minor overall.

### MicroCT

One animal was removed from G3 for a broken distal segment. Bone growth was observed directly upon at least one wire in all samples and was observed upon all four wires in 64% of the samples. This was not significantly different between groups (Figure 5 Left). Non-union was confirmed in all empty defects (groups E3 and E4) and 2/3 of grafted defects (Figure 5 Right). Three of the grafted defects had what appeared to be remnant, unremodeled graft filling the gap but not providing structural bridging. Another three had remodeled graft that failed to unite. Grafted group G3 had significantly higher bone volume (BV) than the two empty groups (p < 0.0001, p = 0.0115, respectively), while group E3 had significantly less total tissue volume (TV) than Groups E4 and G3 (p = 0.0001, p = 0.001, respectively) (Figure 6). As a result, bone volume fraction (BV/TV) and bone mineral density (BMD) followed similar patterns with significantly greater amounts in Group G3 than the two empty groups (p = 0.0007, p = 0.0001, respectively). Tissue mineral density (TMD) was not significantly different between groups (**Error! Reference source not found.**).

**Figure 5.**
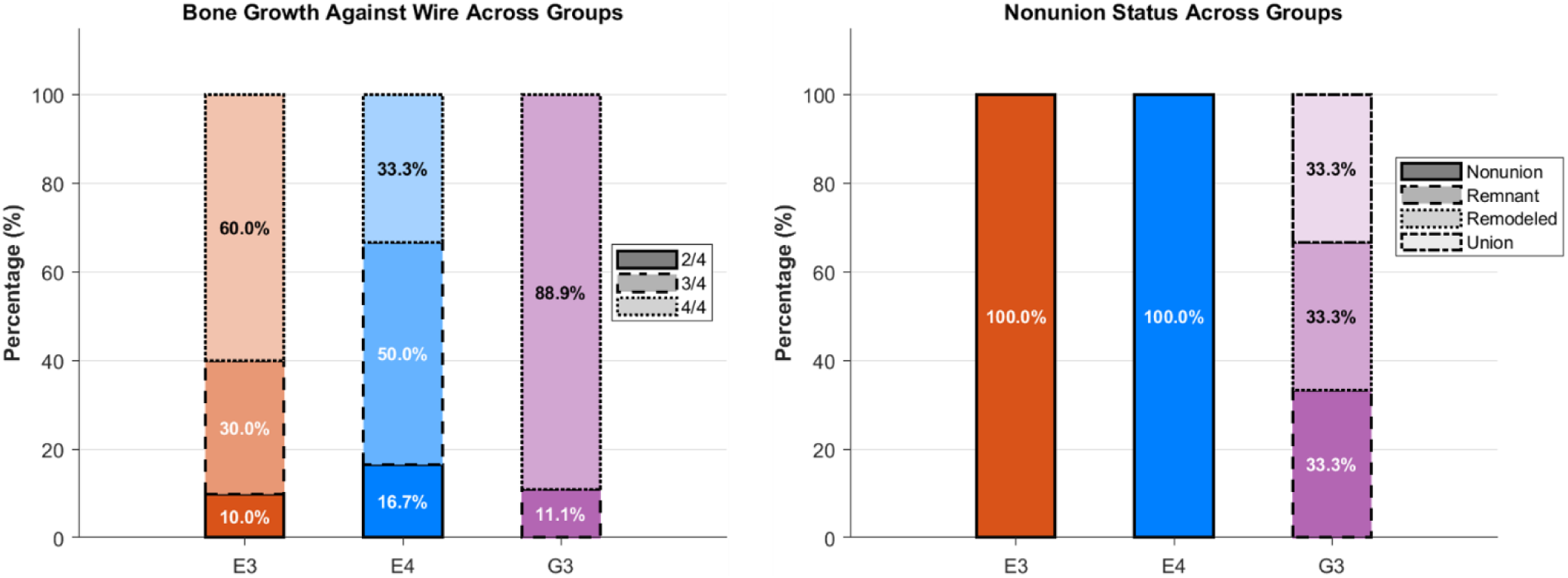
Qualitative microCT results. Bone growth was found on at least 2/4 wires in every sample, regardless of group. Bone growth was present on all four wires in 64% of samples. No samples from E3 or E4 and only 3 from G3 reached union. The remaining G3 samples still contained graft material that was either remnant or remodeled.

**Figure 6.**
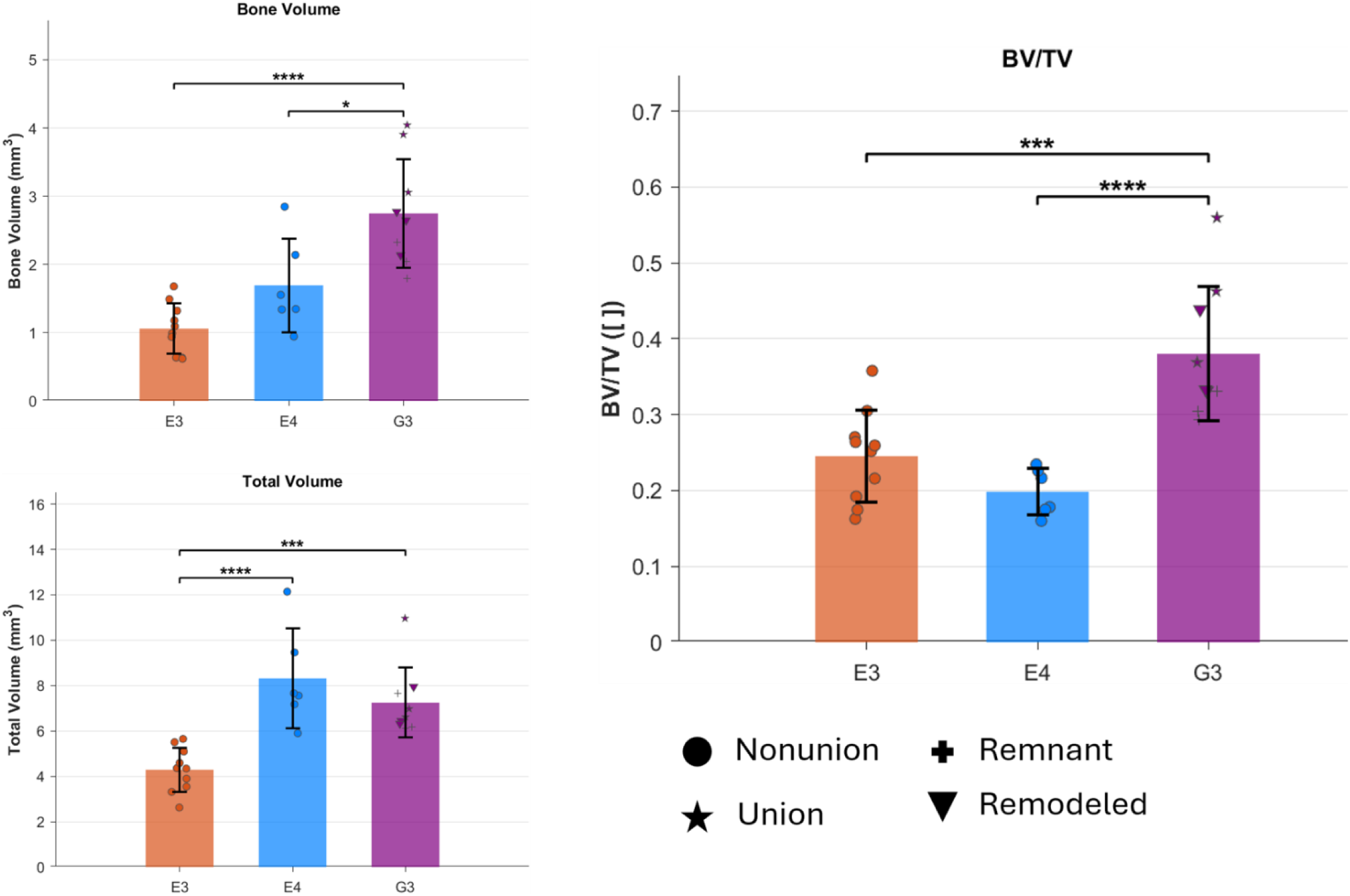
MicroCT bone volume and total volume. G3 had significantly higher bone volume while E3 had lower total volume versus the other groups. BV/TV was higher in G3, but this did not necessarily depend on the united samples as even a remodeled sample had above an average value. P-values: *<0.05, **<0.01, ***<0.001, ****<0.0001

**Figure 7.**
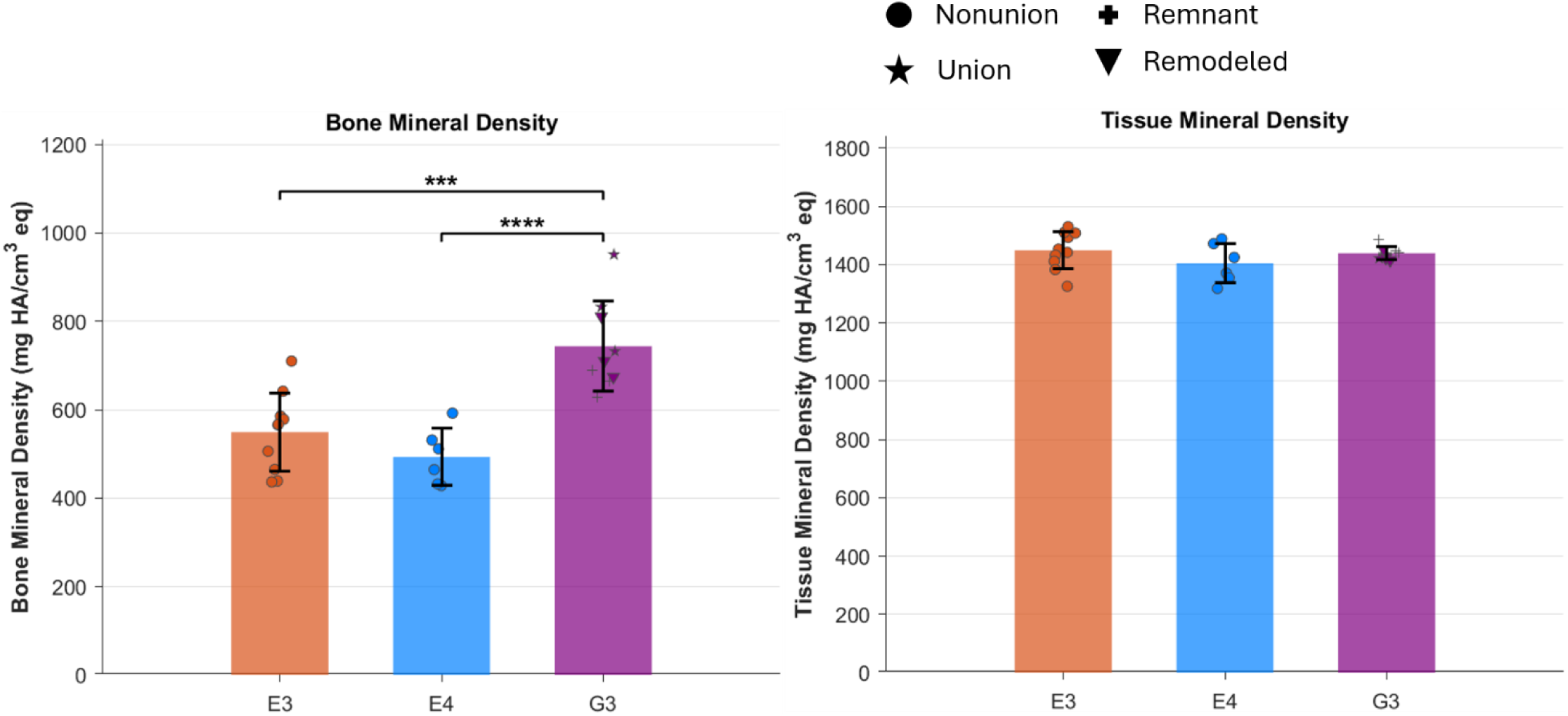
MicroCT bone and tissue mineral density. Bone mineral density was significantly higher in G3 versus both E3 and E4. There were no differences in tissue mineral density. Together this suggests that the bone tissue present is highly similar but different amounts exist in the defect areas, which mirrors the BV/TV results. P-values: *<0.05, **<0.01, ***<0.001, ****<0.0001

## Discussion

This study demonstrates for the first time a viable method to use cerclage wire for plate fixation in a mouse. The study objective was to develop a cost-effective murine CSD model by using cerclage wire for internal plate fixation. Cerclage wire is often used clinically, but as a secondary form of fixation as the wires cannot withstand the internal forces of fracture repair. In small animals, the internal forces are much lower, negating this concern and permitting cerclage wire use as primary fixation at a fraction of the cost. The main challenge in using this method is the possibility of breakage and/or loosening of the wires, leading to complications such as bone resorption or lysis which can impair healing. Our findings support the hypothesis that cerclage wires tied in a modified double-loop fashion can offer stable bicortical fixation with an internal plate.

Some statistical differences were found between groups and over time for the x-ray grading, however, these are functionally minimal to negligible. Plate grades never exceeded a score of 3 (max: 4) while wire grades rarely exceeded a score of 2 (max: 3), indicating that no plates were broken, and both lysis and bone overgrowth were never observed together. Plate scores were not significantly different between the E3 and E4 groups, indicating that the size of the defect does not impact plate stability. In fact, not only did E4 plate scores do not change over time, but E3 scores even slightly improved. Our results reinforce why PEEK is considered the standard radiolucent plating material for rodents, even in more extreme defect scenarios. The wire fixation was shown to *statistically* worsen over time similarly in both empty group; however, fixation timeline in this study far outlasts any previous studies. Very few, if any, studies go past 12 weeks post-op [36–38]. Even with our extended observation period, there were no examples of both bone lysis and overgrowth in the same sample that would indicate on-going or developing instability. The proximal region of the E3 group, which significantly underperformed compared both to the distal wires in the E3 group and the proximal region of the E4 group, was less than half a grade point from comparison wires. Therefore, we do not consider them to be functionally different. These small differences could be due in part to the proximity of this wire to the hip joint as the differing gait pattern of each mouse could contribute to different wear patterns on the proximal most wire. Further, T1500 plated 3 mm defects, which demonstrated exceptional misalignment if not outright plate failure, had similar wire scores across all collected timepoints (Figure S4, S5). This suggests that the wire configuration can withstand exceptional plate material instabilities and still integrate well with bone tissue itself.

MicroCT analysis further supports the overall stability of the wires over time. All samples demonstrated bone growth directly against the wires; 64% of samples had direct growth on all four wires. If the wires were causing on-going instabilities and bone surface damage, bone would not so closely encase them. It is well established from pre-clinical implant studies that on-going micromotion will result in fibrotic encapsulation which appears in microCT as a void area between the implant surface and surrounding mineralized tissues with the area’s width increasing proportionally with instability. In sum, our results suggest satisfactory plate fixation with our wire construct and that defect size does not impact the fixation stability.

Our results also support that 3mm defects utilizing this fixation method will result in non-union if left untreated and limited reconstruction even with grafting. MicroCT analysis confirmed non-union in all empty samples and 69% of grafted samples, reflecting clinical outcomes. Grafted group G3 did have significantly increased BMD, BV/TV, BV, and TV compared to Groups E3 and E4. Superficially this suggests that the grafted group has increased healing, but more critical and rigorous evaluation showed this to not necessarily be the case. Even defects that were categorized as “united” were not without their issues (Figure 8). Sample M341 showed unusual healing patterns as only the medial most part of the defect achieved union while the lateral most part of the defect is mostly empty. Similarly, sample M342 has holes within the united defect. Of the united samples, we feel that only M340 could reasonably bear weight without the aid of fixation. When observing microCT quantitative results, it can be misleading since the three united samples have the highest BV values. However, this does not extend to BV/TV where M342 is less than the average while a remodeled sample that did not achieve union is on par with the other united samples. Without intervention, all human CSDs will result in non-union and the use of a graft alone will not always result in satisfactory bridging, let alone full union [39]. Our results suggest that our murine CSD model is clinically relevant and can be used effectively to detect intervention improvements.

**Figure 8.**
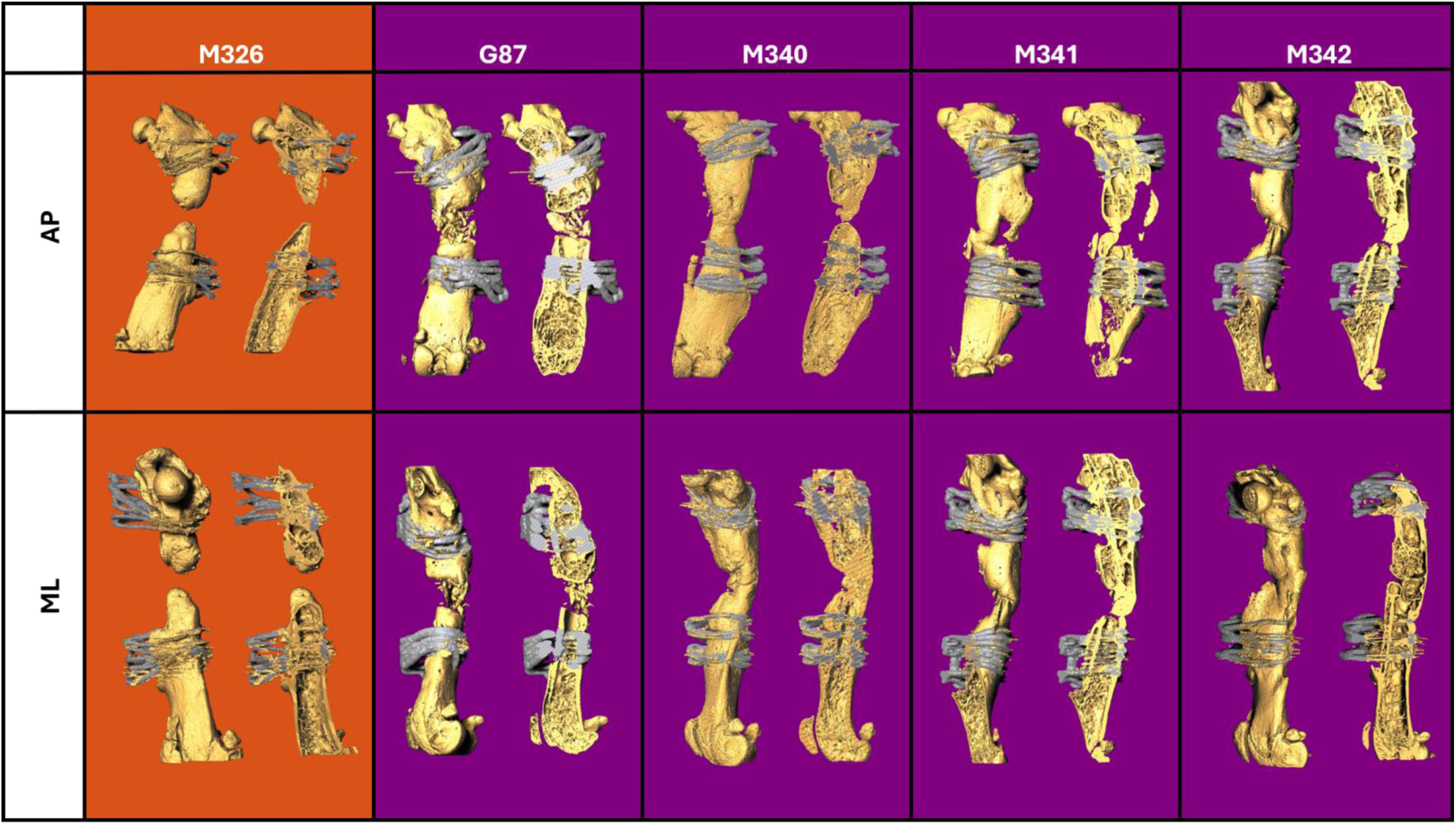
MicroCT reconstructions of representative samples from the empty and grafted groups, as well as the three united samples. Antero- posterior view (top) and medial-lateral view (bottom). Each reconstruction also has a sectional view. M326 (E3) did not receive a graft and consequently failed to unite, instead exhibiting rounded off cortices at the cut ends. G87 (E4) received a graft but did not achieve union; it still has remnant graft material in the defect but is unlikely to unite. M340 (G3) achieved union. M341 (G3) technically united, but only at the medial most part of the defect. M342 (G3) also achieved union but has holes and abnormalities.

As mentioned previously, the use of cerclage wires as primary plate fixation in small animals has only been attempted twice before in a set of companion papers by Burastero *et al.* [31,32]. The purposes of these studies were to combine human mesenchymal stem cells (hMSCs) and BMP-7 with various adjuvants to better treat CSDs in rats. In this model, four cerclage wires were wrapped around both the plate and the whole bone instead of having the wires pass transcortically as well. It is difficult to compare this model to ours as they used rats, there were no empty defect groups, and the efficacy of the wires themselves was not studied. There is not enough information provided in either paper to make definitive *ad hoc* judgements about the wire’s overall performance. However, the provided x-rays at 12 and 16 weeks appear to show adequate fixation was achieved. To our knowledge, this is the most similar study in existence, emphasizing the novelty of the successful use of cerclage wires for internal plate fixation in a murine CSD model.

While our model appears clinically relevant and is cost effective, we also recognize its drawbacks. The main challenge is the learning curve associated with performing the surgery. As discussed in the introduction, a mouse’s small size makes CSD survival surgeries difficult in general, so this is not unique to our model. Any other method of CSD fixation will also be difficult at this scale [25]. Additionally, outcome measurements which require the removal of the implants will be difficult since the wires are so well and complexly integrated into the bone and soft tissues. However, histological analysis should be feasible since the segment of interest (i.e. the defect) can easily be removed from the whole bone after fixation and decalcification for histological processing. MicroCT can be challenging due to beam hardening, but it is achievable when using the correct orientations to minimize any overlapping noise between samples. Alternatively the central area can be removed from the whole fixed bone prior to scanning using precision saws. Mechanical testing, though, could be impossible if evaluating only the regenerate strength is desired. Wire and plate removal for these experiments would likely damage the bone tissue rendering testing impossible or highly inaccurate.

## Limitations

This study is not without limitations. First and primarily, we did not directly compare the efficacy of microscrews and cerclage wire construct strengths ex vivo or in vitro. We felt that direct mechanical comparison would require anatomical loading conditions (i.e. axial loading at the femoral head and condyles), which is extremely difficult to perform correctly and typically requires 10+ specimens to achieve Power.[20] We did not feel the additional animal life was warranted for precise mechanical comparisons at this time. Second, only one tying method was formally tested in this study. Preliminary tests of different tying methods were conducted before deciding on this final form, but these methods’ instability often resulted in early animal removal due to clearly negative effects on the existing bone. Additionally, x-ray grading is an inherently subjective measurement, even when blinded and using standard grading practices. However, it is a commonly used clinical practice, and three highly experienced orthopaedic trauma surgeons from different practices did the grading for this study. So, we employed this method as rigorously as possible within the limitations of the method itself. Similarly, microCT beam hardening prevented quantitative analysis of the peri-wire areas leaving only qualitative wire outcomes possible. While removal of all or part of the wires in preparation for scanning may be possible in the future, we felt that it would distort our findings in this study. Preservation of the *in situ* bone formation was a priority over perfect scanning in these regions.

## Conclusion

CSD intervention is an underpopulated research area, likely due to the financial barriers of using small animal models. With the validation of this affordable murine CSD model, more and increasingly rigorous investigations can now be conducted in transgenic mice, enabling researchers to probe complex biological questions that were previously difficult to address. Our findings demonstrate for the first time that cerclage wires provide an affordable, clinically translatable CSD model that reproduces key clinical outcomes, thereby enabling broader investigation of critical-sized defects and facilitating the development of novel bone repair strategies.

## Supporting information

Supplemental Methods and Results

Supplemental File 1. X-ray Grading Guide

## Acknowledgements

This study was supported by NIH (R21AR080810). We thank Dr. Stacey Meeker for assistance with animal care and surgical refinement. We also thank Joellen Knepfle for her assistance in post-surgical follow-ups and x-ray maintenance.

## References

[1] Schemitsch EH. Size Matters: Defining Critical in Bone Defect Size! J Orthop Trauma 2017;31:S20. 10.1097/BOT.0000000000000978.

[2] Masquelet AC, Fitoussi F, Begue T, Muller GP. [Reconstruction of the long bones by the induced membrane and spongy autograft]. Ann Chir Plast Esthet 2000;45:346–53.

[3] Harris JS, Bemenderfer TB, Wessel AR, Kacena MA. A review of mouse critical size defect models in weight bearing bones. Bone 2013;55:241–7. 10.1016/j.bone.2013.02.002.

[4] Pipitone PS, Rehman S. Management of Traumatic Bone Loss in the Lower Extremity. Orthop Clin North Am 2014;45:469–82. 10.1016/j.ocl.2014.06.008.

[5] Bezstarosti H, Metsemakers WJ, van Lieshout EMM, Voskamp LW, Kortram K, McNally MA, et al. Management of critical-sized bone defects in the treatment of fracture-related infection: a systematic review and pooled analysis. Arch Orthop Trauma Surg 2021;141:1215–30. 10.1007/s00402-020-03525-0.

[6] Huang EE, Zhang N, Ganio EA, Shen H, Li X, Ueno M, et al. Differential dynamics of bone graft transplantation and mesenchymal stem cell therapy during bone defect healing in a murine critical size defect. J Orthop Transl 2022;36:64–74. 10.1016/j.jot.2022.05.010.

[7] Lasanianos NG, Kanakaris NK, Giannoudis PV. Current management of long bone large segmental defects. Orthop Trauma 2010;24:149–63. 10.1016/j.mporth.2009.10.003.

[8] Schmidt AH. Autologous bone graft: Is it still the gold standard? Injury 2021;52:S18–22. 10.1016/j.injury.2021.01.043.

[9] Alford AI, Nicolaou D, Hake M, McBride-Gagyi S. Masquelet’s induced membrane technique: Review of current concepts and future directions. J Orthop Res 2021;39:707–18. 10.1002/jor.24978.

[10] Kaneko Y, Minehara H, Sonobe T, Kameda T, Sekiguchi M, Matsushita T, et al. Differences in macrophage expression in induced membranes by fixation method – Masquelet technique using a mouse’s femur critical-sized bone defect model. Injury 2024;55. 10.1016/j.injury.2023.111135.

[11] Ahmed H, Shakshak M, Trompeter A. A review of the Masquelet technique in the treatment of lower limb critical-size bone defects. Ann R Coll Surg Engl 2023. 10.1308/rcsann.2023.0022.

[12] Fung B, Hoit G, Schemitsch E, Godbout C, Nauth A. The induced membrane technique for the management of long bone defects: a systematic review of patient outcomes and predictive variables. Bone Jt J 2020;102-B:1723–34. 10.1302/0301-620X.102B12.BJJ-2020-1125.R1.

[13] Mauffrey C, Barlow BT, Smith W. Management of Segmental Bone Defects. JAAOS - J Am Acad Orthop Surg 2015;23:143. 10.5435/JAAOS-D-14-00018R1.

[14] Toogood P, Miclau T. Critical-Sized Bone Defects: Sequence and Planning. J Orthop Trauma 2017;31:S23. 10.1097/BOT.0000000000000980.

[15] Muschler GF, Raut VP, Patterson TE, Wenke JC, Hollinger JO. The Design and Use of Animal Models for Translational Research in Bone Tissue Engineering and Regenerative Medicine. Tissue Eng Part B Rev 2010;16:123–45. 10.1089/ten.teb.2009.0658.

[16] Loiselle AE, Zuscik MJ. Skeletal Healing. Primer Metab. Bone Dis. Disord. Miner. Metab., John Wiley & Sons, Ltd; 2018, p. 101–7. 10.1002/9781119266594.ch13.

[17] Kobayashi T, Kronenberg HM. Overview of Skeletal Development. In: Hilton MJ, editor. Skelet. Dev. Repair Methods Protoc., Totowa, NJ: Humana Press; 2014, p. 3–12. 10.1007/978-1-62703-989-5_1.

[18] Gouron R. Surgical technique and indications of the induced membrane procedure in children. Orthop Traumatol Surg Res 2016;102:S133–9. 10.1016/j.otsr.2015.06.027.

[19] Manigrasso MB, O’Connor JP. Characterization of a Closed Femur Fracture Model in Mice. J Orthop Trauma 2004;18:687.

[20] Holstein JH, Garcia P, Histing T, Kristen A, Scheuer C, Menger MD, et al. Advances in the Establishment of Defined Mouse Models for the Study of Fracture Healing and Bone Regeneration. J Orthop Trauma 2009;23:S31. 10.1097/BOT.0b013e31819f27e5.

[21] Garcia P, Holstein JH, Histing T, Burkhardt M, Culemann U, Pizanis A, et al. A new technique for internal fixation of femoral fractures in mice: Impact of stability on fracture healing. J Biomech 2008;41:1689–96. 10.1016/j.jbiomech.2008.03.010.

[22] Histing T, Garcia P, Matthys R, Leidinger M, Holstein JH, Kristen A, et al. An internal locking plate to study intramembranous bone healing in a mouse femur fracture model. J Orthop Res 2010;28:397–402. 10.1002/jor.21008.

[23] Garcia P, Herwerth S, Matthys R, Holstein JH, Histing T, Menger MD, et al. The LockingMouseNail—A New Implant for Standardized Stable Osteosynthesis in Mice. J Surg Res 2011;169:220–6. 10.1016/j.jss.2009.11.713.

[24] Williams JN, Li Y, Valiya Kambrath A, Sankar U. The Generation of Closed Femoral Fractures in Mice: A Model to Study Bone Healing. J Vis Exp JoVE 2018:58122. 10.3791/58122.

[25] Gunderson ZJ, Campbell ZR, McKinley TO, Natoli RM, Kacena MA. A Comprehensive Review of Mouse Diaphyseal Femur Fracture Models. Injury 2020;51:1439–47. 10.1016/j.injury.2020.04.011.

[26] Lenz M, Perren SM, Richards RG, Mückley T, Hofmann GO, Gueorguiev B, et al. Biomechanical performance of different cable and wire cerclage configurations. Int Orthop 2013;37:125–30. 10.1007/s00264-012-1702-7.

27. [27] Stiffler KS. Internal fracture fixation. Clin Tech Small Anim Pract 2004;19:105–13. 10.1053/j.ctsap.2004.09.002.

[28] Angelini A, Battiato C. Past and present of the use of cerclage wires in orthopedics. Eur J Orthop Surg Traumatol 2015;25:623–35. 10.1007/s00590-014-1520-2.

[29] Appleton AT, Mantz GA, Golden-Appleton A, Debenham CT, Sandoval TN, Wang Q, et al. Manual of Wiring Techniques for Orthopedic Surgery. Tech Orthop 2020;35:137. 10.1097/BTO.0000000000000327.

[30] Meyer DC, Ramseier LE, Lajtai G, Nötzli H. A new method for cerclage wire fixation to maximal pre-tension with minimal elongation to failure. Clin Biomech 2003;18:975–80. 10.1016/S0268-0033(03)00181-5.

[31] Burastero G, Scarfì S, Ferraris C, Fresia C, Sessarego N, Fruscione F, et al. The association of human mesenchymal stem cells with BMP-7 improves bone regeneration of critical-size segmental bone defects in athymic rats. Bone 2010;47:117–26. 10.1016/j.bone.2010.03.023.

[32] Burastero G, Sessarego N, Grappiolo G, Castellazzo C, Castello S, Pitto A, et al. Association of ex-vivo expanded human mesenchymal stem cells and rhBMP-7 is highly effective in treating critical femoral defect in rats. J Orthop Traumatol 2007;8:49–54. 10.1007/s10195-007-0163-z.

[33] Katzer A, Marquardt H, Westendorf J, Wening JV, von Foerster G. Polyetheretherketone—cytotoxicity and mutagenicity in vitro. Biomaterials 2002;23:1749–59. 10.1016/S0142-9612(01)00300-3.

[34] Haleem A, Javaid M. Polyether ether ketone (PEEK) and its 3D printed implants applications in medical field: An overview. Clin Epidemiol Glob Health 2019;7:571–7. 10.1016/j.cegh.2019.01.003.

35. Emmer KM, Celeste NA, Bidot WA, Perret-Gentil MI, Malbrue RA. Evaluation of the Sterility of Press’n Seal Cling Film for Use in Rodent Surgery. J Am Assoc Lab Anim Sci JAALAS 2019;58:235–9. 10.30802/AALAS-JAALAS-18-000096.

[36] Liu K, Li D, Huang X, Lv K, Ongodia D, Zhu L, et al. A murine femoral segmental defect model for bone tissue engineering using a novel rigid internal fixation system. J Surg Res 2013;183:493–502. 10.1016/j.jss.2013.02.041.

[37] Zwingenberger S, Niederlohmann E, Vater C, Rammelt S, Matthys R, Bernhardt R, et al. Establishment of a femoral critical-size bone defect model in immunodeficient mice. J Surg Res 2013;181:e7–14. 10.1016/j.jss.2012.06.039.

[38] Clough BH, McCarley MR, Gregory CA. A Simple Critical-sized Femoral Defect Model in Mice. J Vis Exp JoVE 2015:52368. 10.3791/52368.

[39] Findeisen S, Schwilk M, Haubruck P, Ferbert T, Helbig L, Miska M, et al. Matched-Pair Analysis: Large-Sized Defects in Surgery of Lower Limb Nonunions. J Clin Med 2023;12:4239. 10.3390/jcm12134239.

