## Supplemental Methods and Results for "Cerclage Wire as an Affordable Alternative for Internal Fixation in Murine Critical Sized Defect Models"

***Supplementary Methods & Results***

*Surgical Details*

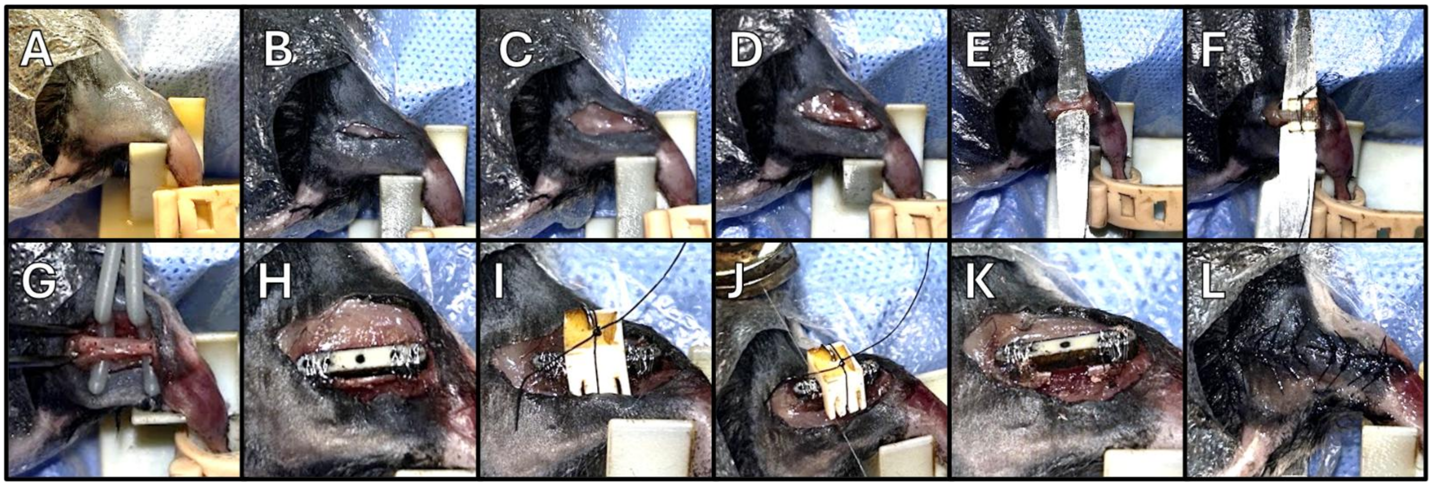


***Figure S1.*** *Surgical Procedure.* ***A.*** *Right hind limb prepped and secured in surgical stand.* ***B.*** *Incision made from just proximal of trochanteric ridge to femoral condyles.* ***C.*** *Fascia bluntly dissected.* ***D.*** *Muscle bluntly dissected to exposed femur.* ***E.*** *Flat metal plate is slid over the muscles and under the bone to protect soft tissue during drilling.* ***F.*** *Drill guide is attached to the femur using suture and four 0.4mm holes are drilled.* ***G.*** *Drill holes.* ***H.*** *Plate is secured to femur using four wires.* ***I.*** *Saw guide is lined up with the middle hole of the plate and secured using suture.* ***J.*** *Two cuts are made to form a defect.* ***K.*** *3mm defect.* ***L.*** *Incision is closed using 5-0 monofilament in interrupted surgeon's knots.*

*Tough 1500 as a Mechanically Compromised Plate Material*

Tough 1500 is a 3D resin from FormLabs that is known to be biologically compatible and autoclavable. Additionally, plates were readily available for fabrication from the Medical Modeling, Materials and Manufacturing (M4) Lab at The Ohio State University and could be printed with complex geometries for a fraction of what a similarly complex PEEK plate would cost (Figure S2). However, since very little real application had been done with T1500 at the start of this study, we wanted to ensure its viability as an *in vivo* implant.

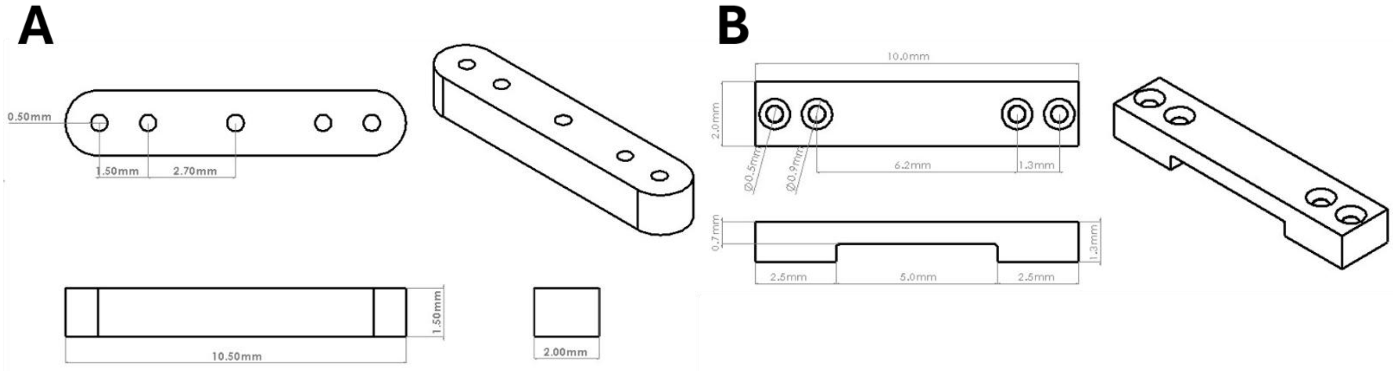


***Figure S2.*** *Plate Dimensions.* ***A.*** *PEEK plate dimensions. PEEK plates were laser cut from a 1.5 mm thick sheet of PEEK by the 3D Printing Laboratory in the Department of Mechanical and Aerospace Engineering.* ***B.*** *Tough 1500 plates. T1500 plates were resin printed by the M4 Lab at The Ohio State University.*

Tough 1500 and PEEK plates were exposed to either phosphate buffer solution (PBS) or air at either room temperature (20°C) or body temperature (40°) to replicate the murine *in vivo* environment. Plates were kept in this environment for 24 or 48 hours (n = 4/group; N = 32). Plates were then subjected to 3-point bend testing to failure (preload of 3N, 0.1 mm/s displacement, Instron Elecropuls E3000). Resulting force-displacement data was converted to stress-strain using a custom matlab program (see table Figure S3B for input parameters). Resulting data was analyzed using a multivariate ANOVA with factors of medium, temperature, and time. Time and temperature were found to not be significant. Thus they were pooled to increase power, and a second MANOVA was performed (factors: material, medium). We further blocked by material if a significant covariation was found between the factors (Figure S3C). T1500 performed much worse than PEEK, regardless of medium (Figure S3A). This was expected as PEEK is known to have superior mechanical properties. Interestingly, T1500 was compromised in PBS versus air, indicating that these plates would most likely be compromised *in vivo* due to fluid absorption. Conversely, PEEK was unaffected by PBS exposure, indicating stability *in vivo*.

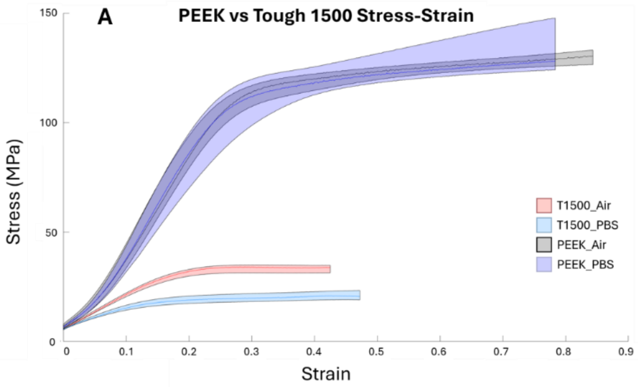


**B**

| Material | Cross Sectional Area ($mm^2$) | Gauge Length (mm) | Distance from Center of Mass (mm) | Moment of Inertia ($mm^4$) |
|---|---|---|---|---|
| PEEK | 3 | 3.9 | 0.75 | 0.4215 |
| T1500 | 2 | 3.9 | 0.5 | 0.25 |

**C**

| Parameter | Material | Medium | Covariation | Cov. PEEK | Cov. T1500 |
|---|---|---|---|---|---|
| Stiffness | **<0.0001** | **0.0131** | 0.6822 | - | - |
| Elasticity | **<0.0001** | **0.0018** | 0.7483 | - | - |
| Rigidity | **<0.0001** | **0.0131** | 0.6822 | - | - |
| Toughness (S-S (J)) | **<0.0001** | 0.1507 | **0.0318** | 0.0823 | **<0.0001** |
| Yield Strain | **<0.0001** | **0.0304** | **0.003** | 0.6417 | **<0.0001** |
| Yield Stress | **<0.0001** | **<0.0001** | 0.2276 | - | - |
| Ultimate Strain | **<0.0001** | **0.0109** | 0.9088 | - | - |
| Ultimate Stress | **<0.0001** | 0.2878 | **0.0001** | **0.0157** | **<0.0001** |
| Break Strain | **<0.0001** | 0.1669 | 0.086 | - | - |
| Break Stress | **<0.0001** | 0.2506 | **0.0002** | **0.016** | **<0.0001** |

***Figure S3.*** *Mechanical testing results.* ***A.*** *Stress-strain curve for each group with standard deviation as cloud. PEEK outperformed T1500 and was not impacted by exposure to PBS while T1500 was.* ***B.*** *Three-point bend input parameters.* ***C.*** *P-values for statistical tests.*

The same survival surgeries were performed on 13 animals to implant a T1500 plate and create a 3mm femoral defect (Group T3). Only one animal out of 13 made it to the 20 week timepoint, all others were removed early for broken plates. When included in the blinded x-ray grading (same statistical methods as PEEK only data shown in the man manisrcipt), T3 plates scored significantly worse than both E3 and E4 groups ($p < 0.001$, $p = 0.008$, respectively) and significantly degraded over time ($p < 0.0001$), supporting the hypothesis formed from the mechanical testing that these plates would be unstable *in vivo* (Figure S4).

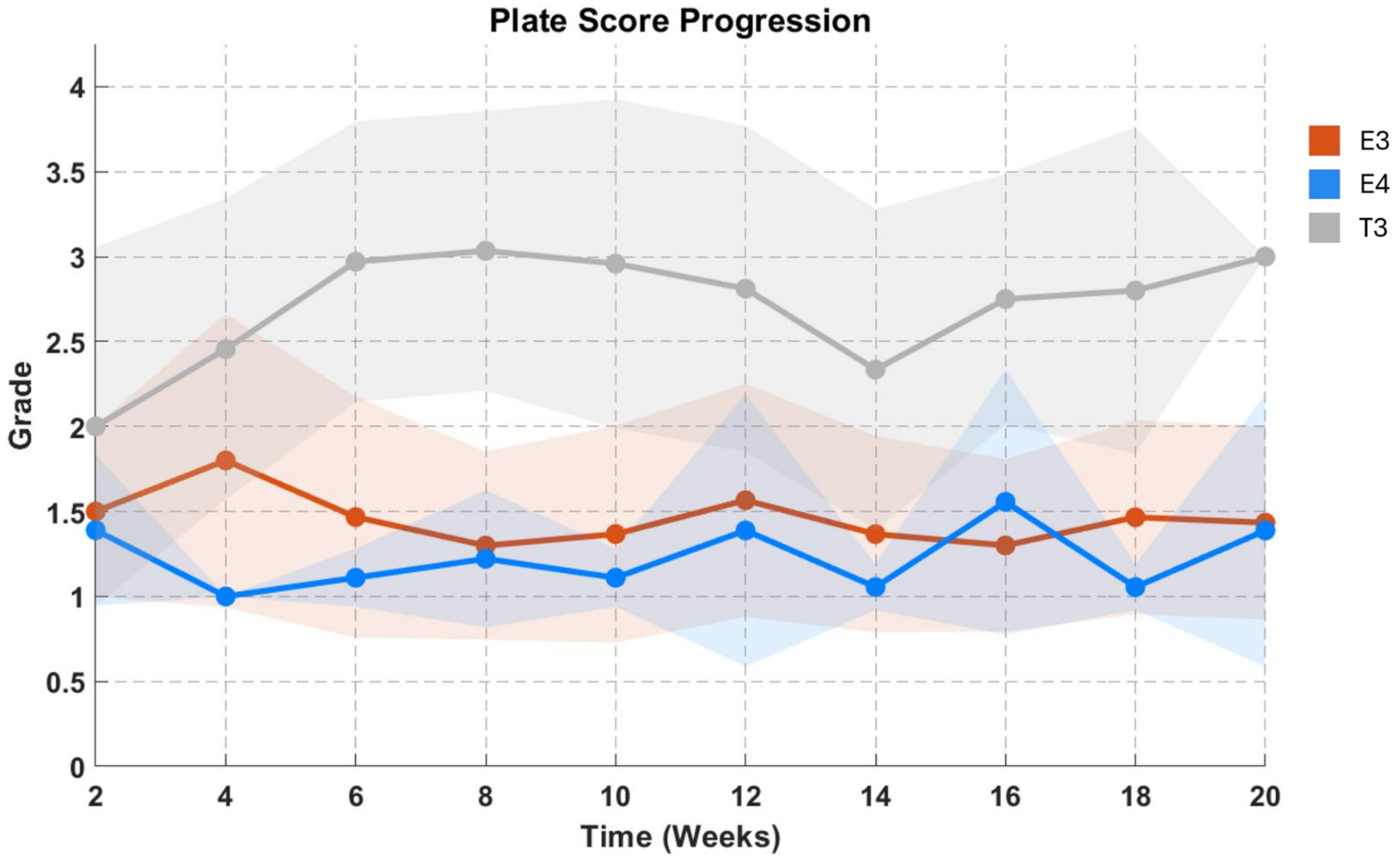


***Figure S4****. Plate score progression including T3. T3 scored significantly worse than E3 and E4 and significantly degraded over time, indicating that T1500 was not stable in vivo.*

The longitudinal worsening rate of T3 wires was significantly worse than that of E3 and E4 ($p = 0.0023$, $p = 0.0012$, respectively). In the proximal most and proximal middle regions, T3 wires were not significantly different from E4, but did score better than E3 ($p = 0.0009$, 0.0071, respectively). There were no significant differences in the distal segment. Additionally, much like the E4 group, there were no significant regional differences in wire scores for T3 (Figure S5).

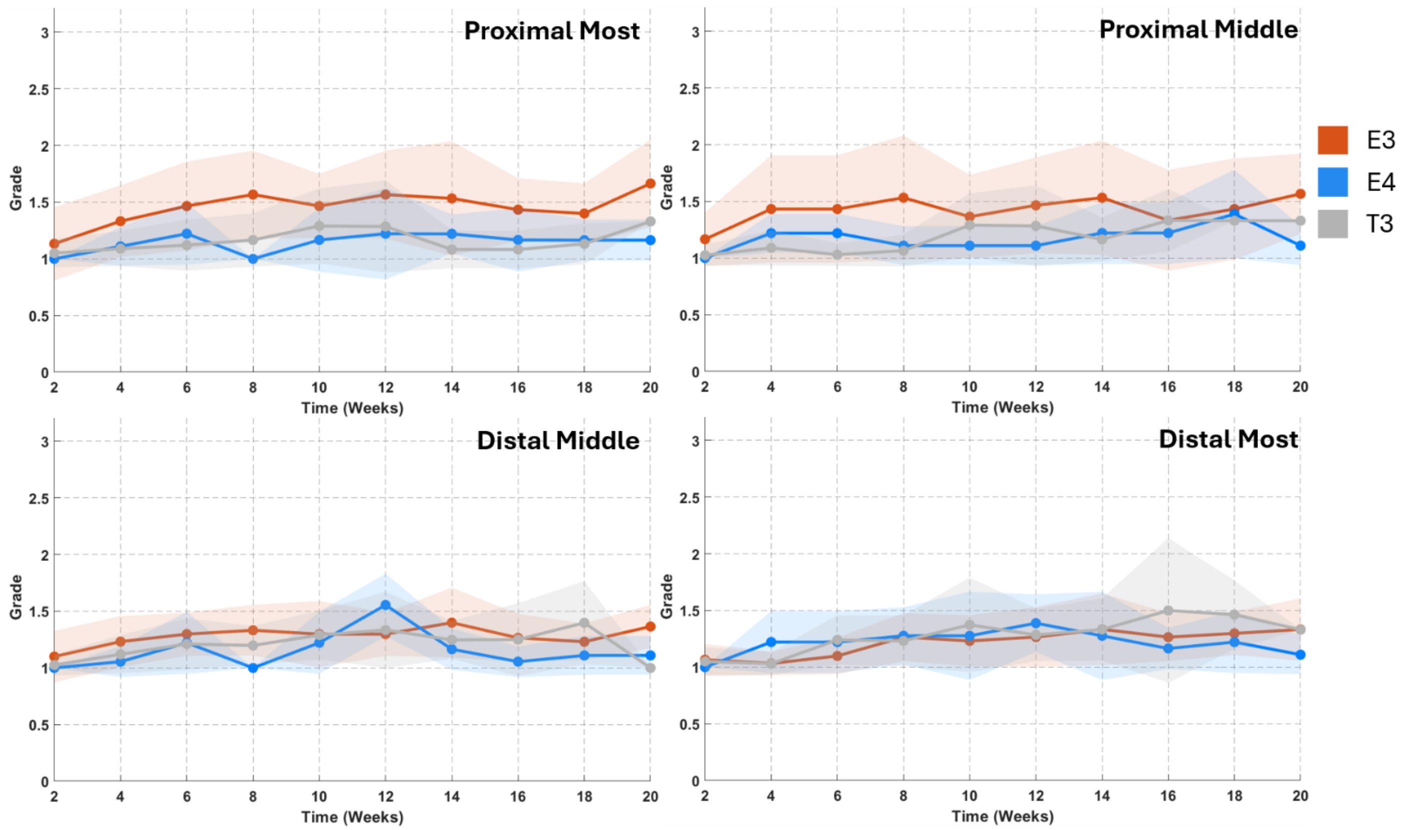


***Figure S5***. *Wire score progression including T3. Longitudinal rate of worsening was significantly worse in T3. The proximal most and proximal middle regions were not different between T3 and E4; T3 also outperformed E3 at these regions.*

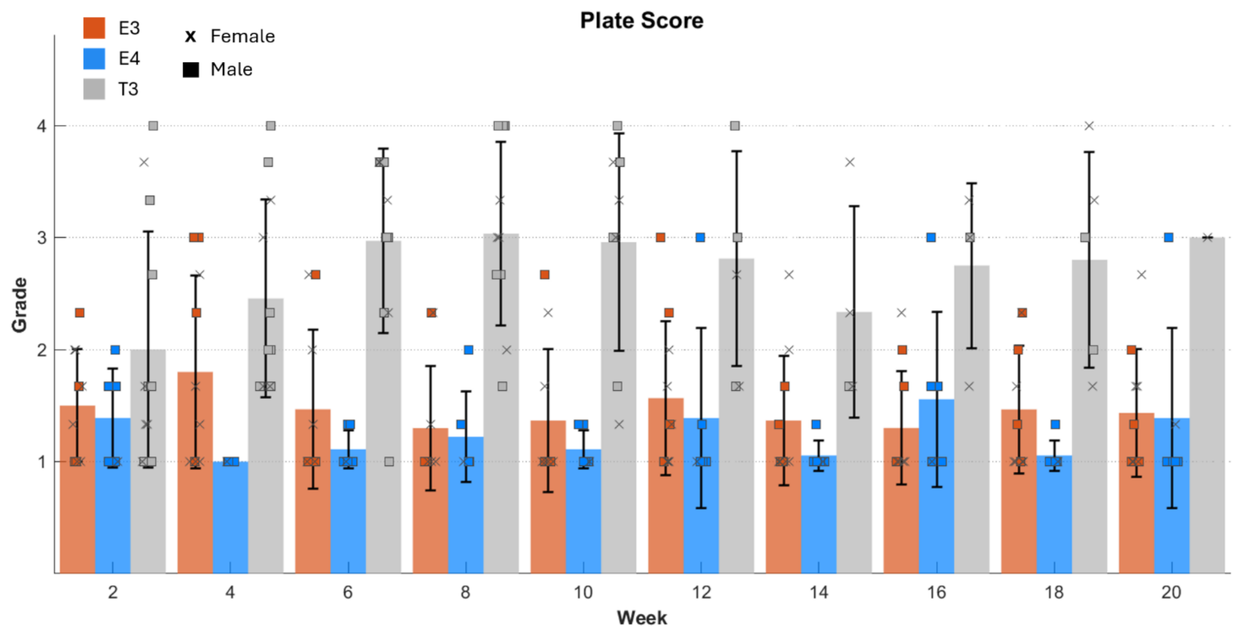


***Figure S6.*** *Plate score progression with sex and individual distribution.*

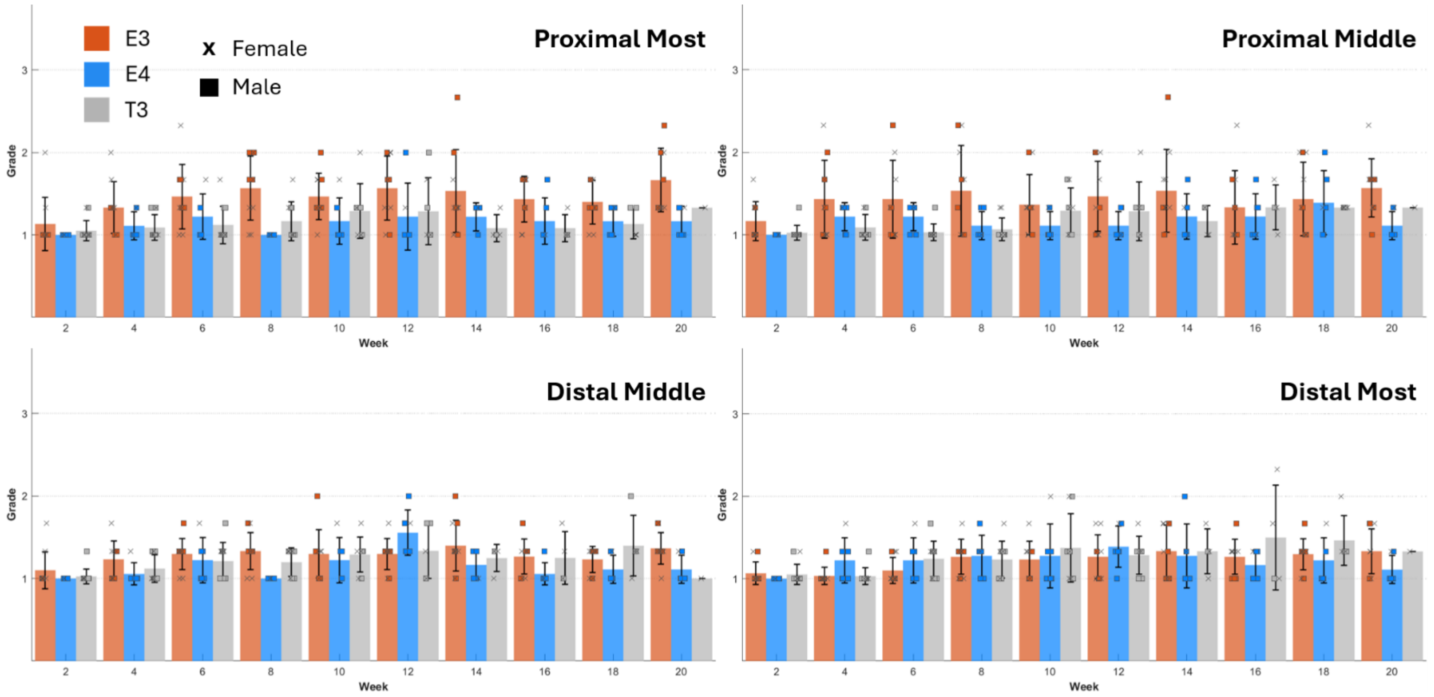


*Figure S7.* *Wire score progression with sex and individual distribution.*

### *All MicroCT reconstructions for PEEK samples*

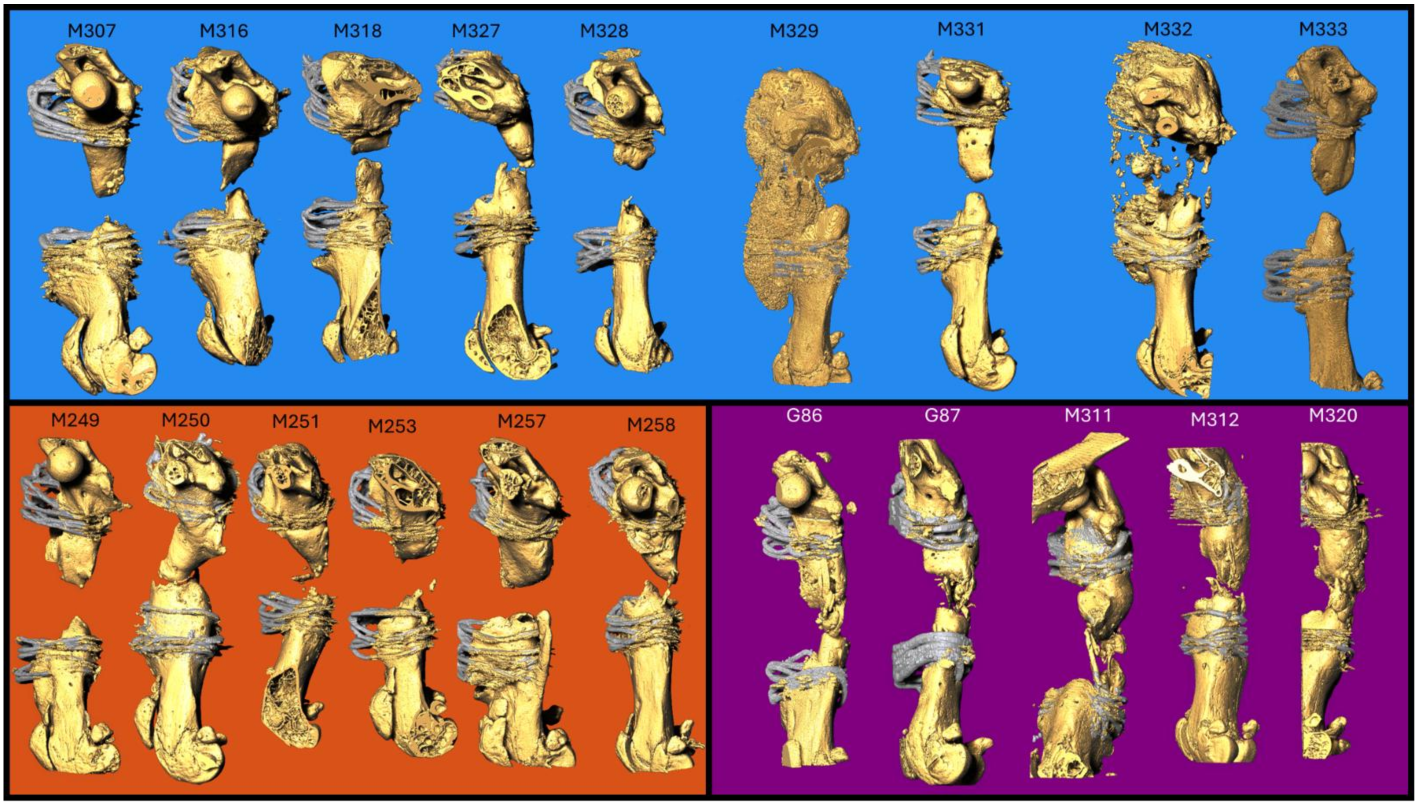


*Figure S8.* *MicroCT reconstructions of all samples at the ML view. All empty samples resulted in the rounding off the cut ends. M329 and M332 in group E3 had a large amount of bone growth onto the plate. None of the G3 groups exhibited here reached union.*
