## Supplemental File 1. X-ray Grading Guide for "Cerclage Wire as an Affordable Alternative for Internal Fixation in Murine Critical Sized Defect Models"

### General Orientation

Each sample has a pair of xrays, each of a different view of the same animal at the same time point. The left image will be looking at the flat surface of the plate while the right image looks at the anterior side of the plate. Given that the plate is not radio-opaque, you will not actually be able to see it, but you can gather its position based on the wires and the knots tied at the top of the plate. For example, below is an xray of a mouse without a critical defect. Looking at the left image, we don't see a large amount of space between the wire loops and the bone since we are looking at the top view of the plate. On the right, however, there is clearly a gap where the plate is given that we are now looking at a side view (indicated by the outlined box).

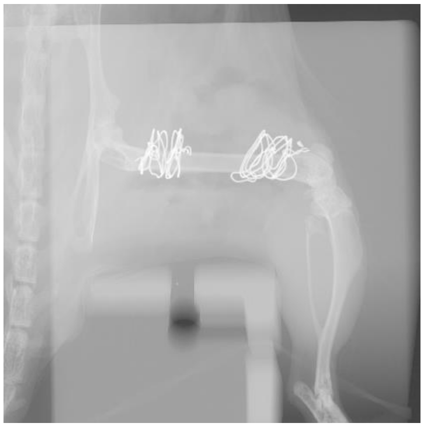

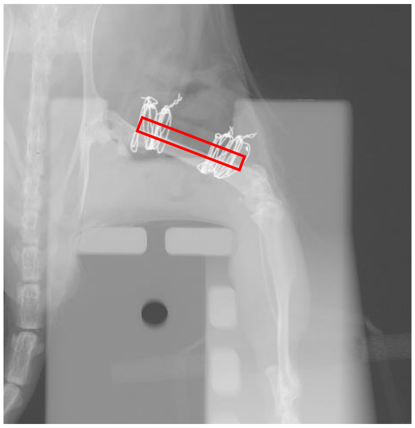

These angles are achieved by adjusting the position of the foot, as seen below.

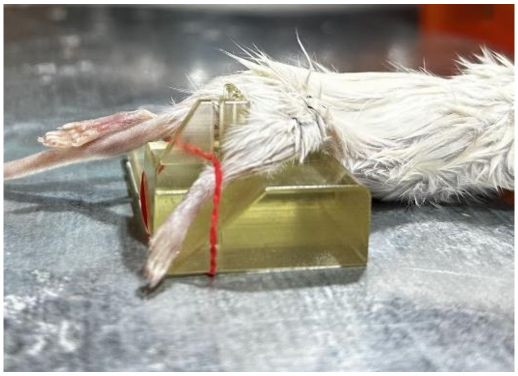

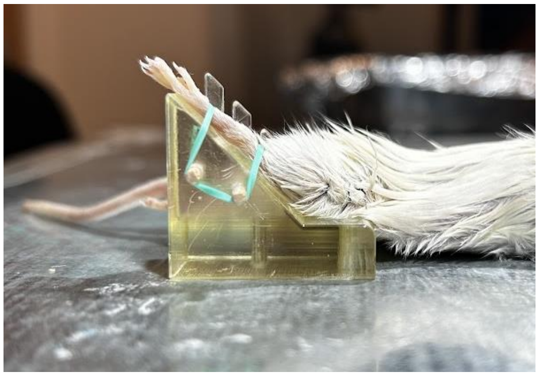

Plate Grading

To grade the status of the plate, use the proximal segment of the femur as the horizontal origin. Any movement of the distal segment relative to the proximal segment can be seen as bending. There are different tiers of bending, as seen below. If the distal segment still looks in line with the proximal, classify it as a 1 for no bend. If there is slight movement in either direction within the yellow lines, rate it a 2 for slight bend. If above the yellow lines, rate it a 3 for major bend. This bend will be easiest to see in this view.

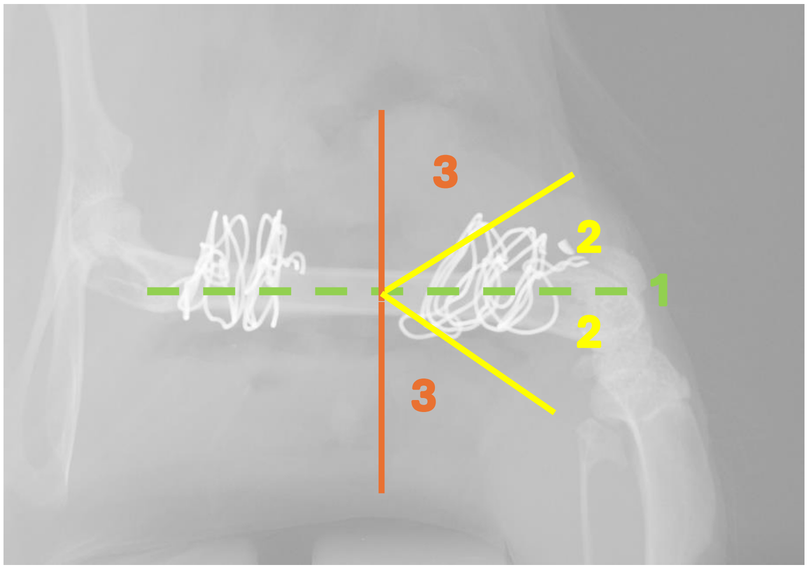


The last tier, indicated by a 4, means that the plate is broken. This can be seen by looking between the views. The proximal segment does not appear to rotate with the distal, this indicates that the plate has broken. Below are the orientations that could indicate a broken plate. Solid green indicates bone in the correct position for that view, solid red indicates bone in the incorrect position for that view, orange represents the wires, the plate is represented by a black outline, and the blue arrows indicate the axis of rotation for the distal segment relative to the proximal.

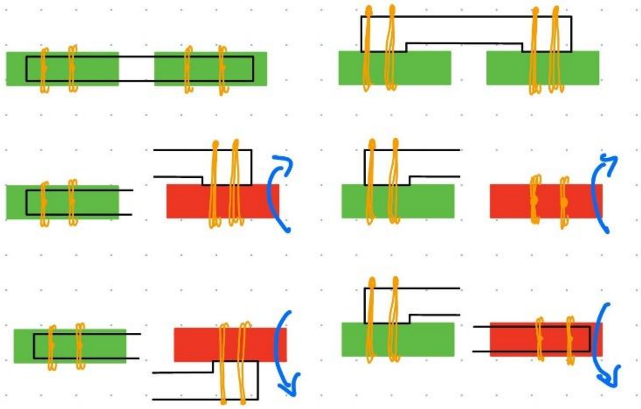

| Plate Grading |
| --- |
| 1 → no bend |
| 2 → slight bend |
| 3 → major bend |
| 4 → broken |

Wire Hole Grading

To grade the security of the wires, look for two indicators: lysis and/or new bone growth around the wires. Lysis is indicated by the widening of the hole through the bone, such as seen below. The more proximal holes are obviously quite larger than the distal holes, despite having been made with the same drill bit. This largening means that the wires are less stable and have less purchase on the bone.

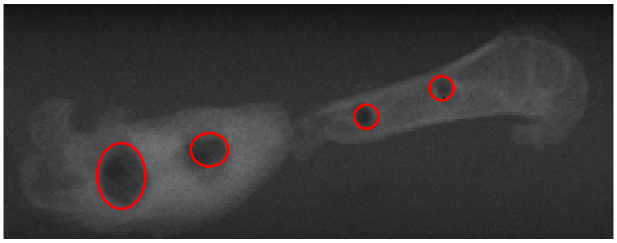

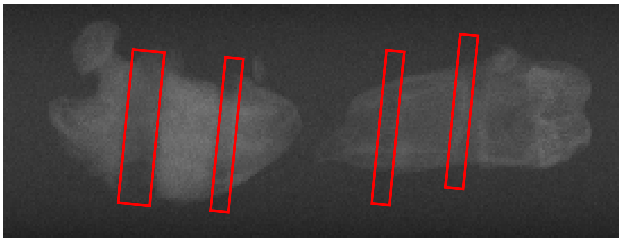

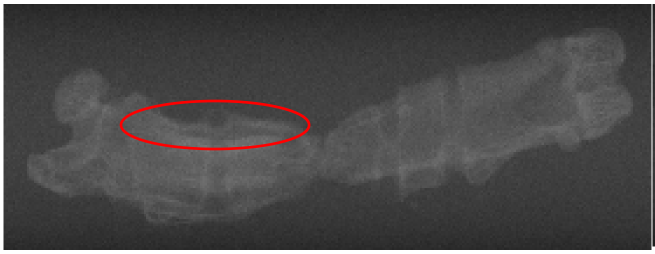

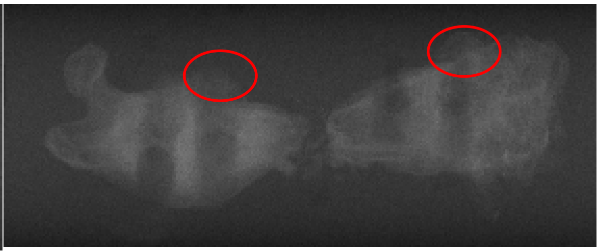

New bone growth is indicated by the presence of a faint outline outside of the original calcified bone. Since the bone is quickly trying to repair itself, it will make emergency bone that is not as mineralized as the original, hence the faintness in xray. There may be some layered bone growth, outlined on the left, which is normal in the repair process. However, bulging bone growth, outlined on the right, is indicative  of instability.

Please rate each of the holes as their own entity, start with the proximal most hole near the hip and ending with the distal most hole near the knee. If the hole shows neither lysis nor new bone growth, rate it as a 1. If it shows only one of these signs, rate it as a 2. If it shows as both, rate it as a 3.

| Wire Hole Grading Scale |
| --- |
| 1 → no lysis or new bone growth |
| 2 → either lysis or new bone growth |
| 3 → lysis and new bone growth |

### Summary

As a summary, each pair of xrays should have **5 total ratings**: 1 for the plate status and 1 for the status of each of the 4 wire holes. You may go back and change your rating at a later date, but do not communicate with anyone else participating in these ratings. Feel free to reach out if you need any further clarification of the guidelines. Once you have finalized your ratings, please let me know.
